# Towards a reliable, automated method of individual alpha frequency (IAF) quantification

**DOI:** 10.1101/176792

**Authors:** Andrew W. Corcoran, Phillip M. Alday, Matthias Schlesewsky, Ina Bornkessel-Schlesewsky

**Affiliations:** Cognition and Philosophy Laboratory, Monash University, Melbourne, Australia.; Centre for Cognitive and Systems Neuroscience, University of South Australia, Adelaide, Australia.; Max Planck Institute for Psycholinguistics, Nijmegen, 6500 AH, The Netherlands.

**Keywords:** Alpha Rhythm, EEG, Oscillation/Time Frequency Analyses, Savitzky-Golay Filter, Individual Alpha Frequency

## Abstract

Individual alpha frequency (IAF) is a promising electrophysiological marker of interindividual differences in cognitive function. IAF has been linked with trait-like differences in information processing and general intelligence, and provides an empirical basis for the definition of individualised frequency bands. Despite its widespread application, however, there is little consensus on the optimal method for estimating IAF, and many common approaches are prone to bias and inconsistency. Here, we describe an automated strategy for deriving two of the most prevalent IAF estimators in the literature: peak alpha frequency (PAF) and centre of gravity (CoG). These indices are calculated from resting-state power spectra that have been smoothed using a Savitzky-Golay filter (SGF). We evaluate the performance characteristics of this analysis procedure in both empirical and simulated EEG datasets. Applying the SGF technique to resting-state data from *n* = 63 healthy adults furnished 61 PAF, and 62 CoG estimates. The statistical properties of these estimates were consistent with previous reports. Simulation analyses revealed that the SGF routine was able to reliably extract target alpha components, even under relatively noisy spectral conditions. The routine consistently outperformed a simpler method of automated peak detection that did not involve spectral smoothing. The SGF technique is fast, open-source, and available in two popular programming languages (MATLAB and Python), and thus can easily be integrated within the most popular M/EEG toolsets (EEGLAB, FieldTrip and MNE-Python). As such, it affords a convenient tool for improving the reliability and replicability of future IAF-related research.

## 1 Introduction

Alpha is the dominant rhythm in the human EEG, and its importance for cognitive processing has been recognised since Hans Berger’s seminal work in the early 20th century (cf. Adrian & Matthews, 1934; Berger, 1929). Interindividual differences in the predominant frequency of alpha-band oscillations (i.e. individual alpha frequency; IAF) have been linked with variability in cognitive performance since the 1930s (for a more recent review, see Klimesch, 1999; see Vogel & Broverman, 1964). More recent research has revealed that IAF predicts performance on a variety of perceptual (e.g., Cecere, Rees, & Romei, 2015; Samaha & Postle, 2015) and cognitive (e.g., Bornkessel, Fiebach, Friederici, & Schlesewsky, 2004; Klimesch, Doppelmayr, & Hanslmayr, 2006) tasks. Individuals with a low IAF process information more slowly (Klimesch, Doppelmayr, Schimke, & Pachinger, 1996; Surwillo, 1961, 1963), and show reduced performance on memory tasks (Klimesch, 1999) and general intelligence measures (*g*; Grandy et al., 2013a), in comparison to their high-IAF counterparts. IAF is a trait-like characteristic of the human EEG (Grandy et al., 2013b), which shows high heritability (Lykken, Tellegen, & Thorkelson, 1974; Malone et al., 2014; Smit, Wright, Hansell, Geffen, & Martin, 2006) and test-retest reliability (Gasser, Bächer, & Steinberg, 1985; Kondacs & Szabo, 1999; Näpflin, Wildi, & Sarnthein, 2007). However, IAF tends to decrease with age from young adulthood onwards (Chiang, Rennie, Robinson, Albada, & Kerr, 2011; Köpruner, Pfurtscheller, & Auer, 1984), hence lifelong changes in IAF accompany the decline of many cognitive abilities in older adulthood (e.g. Hedden & Gabrieli, 2004; Salthouse, 2011). Taken together, this evidence highlights the utility of the IAF as a neurophysiological marker of general brain functioning (Grandy et al., 2013a, 2013b).

In addition to quantifying individual differences in the properties of the dominant alpha rhythm, IAF can also be used to derive individualised estimates of the canonical frequency bands beyond alpha (Klimesch, 2012). Such empirically-driven approaches to frequency band definition have been proposed to sharpen the precision of frequency-domain analyses more generally (Klimesch, 2012). Indeed, using the IAF to distinguish subregions of the alpha band has revealed functional dissociations between lower-and higher-frequency alpha-rhythms (e.g., Klimesch, 1997). However, despite the potential advantages of deploying the IAF as a reference point for various kinds of individualised spectral analysis, no clear consensus on the optimal method for quantifying IAF currently exists. This paper thus sets out to develop a rigorous, automated strategy for estimating two of the most widely reported indices of IAF: peak alpha frequency (PAF) and alpha frequency centre of gravity (CoG). We begin by briefly describing some of the most common strategies for extracting these estimators, and their attendant problems.

### 1.1 Peak alpha frequency

IAF estimation typically depends on the delineation of a singular, prominent spectral peak within the alpha bandwidth (standardly defined as 8-13 Hz; Noachtar et al., 2004). In many cases, the PAF can be easily discerned upon visual inspection of the power spectral density (PSD) from eyes-closed resting-state EEG recorded over parieto-occipital sites. However, this strategy is complicated by the presence of two (or more) alpha-band peaks (so-called “split-peaks”; Chiang et al., 2011), or the lack of any obvious deviation from the characteristic 1*/f*-like scaling of background M/EEG spectral activity (the “inverse power-law”; Pritchard, 1992). Under such circumstances, subjective PAF estimation may be prone to bias and inconsistency (Chiang et al., 2008), thus posing a significant challenge to replicability. While conservative approaches to PAF identification in the context of ambiguous spectral conditions may help reduce bias, this may result in high rates of attrition (see for e.g., Bornkessel-Schlesewsky et al., 2015).

One approach for improving the objectivity, replicability, and (for larger datasets) practicality of PAF estimation is to implement an automated peak-detection algorithm. While this strategy does not solve the basic problem of deciding the criteria by which valid PAF estimates are discriminated from split-peaks or spurious background fluctuations, it at least applies such criteria consistently across all subjects. Simple algorithms may however introduce new sources of bias. For instance, a basic routine that searches for local maxima within the alpha band may arbitrarily assign the PAF to the lower bound of the search window in the absence of any notable deviation from the inverse-power law (since the highest power estimate will be the supremum found at the lowest frequency bin spanned by the window). A more sophisticated approach such as the first-derivative test (in which the first derivative of the PSD is searched for downward going zero crossings; cf. Grandy et al., 2013b) avoids this problem, but is still incapable of distinguishing substantive peaks from split-peaks or arbitrarily small deviations from background spectral activity. Such routines may therefore be too liberal with regard to the spectral features they classify as alpha peaks.

### 1.2 Alpha-band centre of gravity and reactivity

The alpha mean or CoG frequency (Klimesch, Schimke, Ladurner, & Pfurtscheller, 1990) has been proposed as an alternative method of IAF estimation that circumvents some of the difficulties posed by the absence of a dominant alpha peak (Klimesch, 1997; Klimesch, Schimke, & Pfurtscheller, 1993). This estimator computes a weighted average of the power contained within the alpha-band, thus rendering a summary measure that is sensitive to the spectral distribution of alpha components. Given that the span and location of alpha-rhythm activity vary across individuals (Bazanova & Vernon, 2014), Klimesch and colleagues (1990) recommended computing the CoG using bespoke frequency windows designed to capture such variation. However, the definition of such individualised alpha-band windows (IAWs) poses a nontrivial challenge, and may rely on subjective assessments or arbitrary criteria (Bazanova & Vernon, 2014). One principled solution to this problem is to derive the IAW from reactivity-based contrasts between two conditions (pre-vs. peri-stimulus presentation, Goljahani et al., 2012; e.g., eyes-closed vs. eyes-open resting-states, Klimesch, 1999). This approach is not immune to bias, however, since alpha rhythms are not always substantially attenuated by opening the eyes (Gaál, Boha, Stam, & Molnár, 2010; Kreitman & Shaw, 1965), and may only be partially attenuated (e.g., Klimesch et al., 2006) – or even *enhanced* (e.g., Rihs, Michel, & Thut, 2007) – during experimental tasks.

### 1.3 Curve-fitting approaches to alpha-rhythm quantification

One promising approach to spectral peak quantification that avoids many of the issues highlighted above applies iterative curve-fitting techniques to parameterise the statistical properties of the PSD (e.g., Chiang et al., 2008; Lodder & Putten, 2011). The practical utility of such methods is clearly apparent from their application to large *n* datasets (Albada & Robinson, 2013; e.g., Chiang et al., 2011), while comparison of Lodder and van Putten’s (2011) algorithm with human scorers revealed a high degree of estimator agreement. It is puzzling then why such methods have not been taken up more broadly within the IAF literature (cf. Haegens, Cousijn, Wallis, Harrison, & Nobre, 2014, for a notable exception). One possibility is that investigators are generally unaware of these approaches, given that they have mostly been applied in the context of spectral modeling rather than IAF research (nor Bazanova and Vernon, 2014, mention the existence of such methods in their reviews of IAF estimation techniques; indeed, neither Goljahani et al., 2012). Alternatively, investigators may be put off by the perceived burden involved in accessing these programmes (which we have not been able to locate publically) and integrating them within existing analysis pipelines (which may not be compatible with such algorithms). We suggest then that one of the critical steps towards achieving a more widespread adoption of automated IAF estimators is to make these tools openly available in formats that can be easily assimilated within popular methods of M/EEG analysis.

### 1.4 Aims of the present study

In sum, common methodological approaches to IAF estimation are either (1) time-consuming and vulnerable to inconsistencies arising from subjective evaluation, or (2) at risk of producing spurious or biased estimates under certain plausible spectral conditions. More recent innovations that address these problems via the application of sophisticated curve-fitting algorithms have so far found limited uptake within the broader IAF literature, perhaps on account of practical barriers pertaining to software access and implementation. Consequently, we sought to develop an automated method of alpha-band quantification that provides fast, reliable, and easily replicated estimates of the resting-state IAF in two major programming languages: MATLAB^®^ (The MathWorks, Inc., Natick, MA, USA) and Python™. This goal is consistent with recent proposals to make the analysis of electrophysiological data as open, transparent, and amenable to replication as possible (Cohen, 2017).

## 2 Method

Our approach aims to emulate Klimesch and colleagues’ (1990) original attempt to characterise individual profiles of resting-state alpha-band activity by means of a relatively simple, non-parametric curve-fitting technique; the Savitzky-Golay filter (SGF). The basic strategy runs as follows: First, we extract PSD estimates from preprocessed, fast Fourier-transformed EEG signals. Second, we apply the SGF to smooth the PSD function and estimate its first-and second-order derivatives. Third, these derivatives are analysed for evidence of a distinct spectral peak within the alpha band region. Finally, the first derivative of the PSD is reanalysed to locate the bounds of the IAW, from which the CoG is estimated. Our main focus here will be to assess the efficacy of this approach in the context of both empirical and simulated data. For a more rigorous account of the calculations implemented in the algorithm, see Appendix.

### 2.1 Savitzky-Golay smoothing and differentiation

The SGF is a least-squares polynomial curve-fitting procedure specifically designed to aid the detection of spectral peaks amidst noisy conditions (Savitzky & Golay, 1964). The major advantage of the SGF in this regard is its ability to smooth peaks while preserving their height, width, position, and CoG (Schafer, 2011; see Ziegler, 1981). Consequently, we propose using the SGF in order to attenuate random fluctuations in the PSD (and thus improve signal-to-noise ratio; SNR) without substantially distorting the spectral parameters of interest in IAF analysis. Eliminating such fluctuations should reduce the number of spurious local optima in the derivatives of the PSD, thus improving the overall accuracy and reliability of the first-derivative test. Conveniently, SGFs constitute optimal (or near optimal) differentiators (Luo, Ying, He, & Bai, 2005), and hence can be deployed to estimate both the smoothed PSD and its derivatives simultaneously.

### 2.2 Implementation

All functions developed in order to conduct the analyses reported here are open-source and available (along with sample datasets and simulation materials) from https://github.com/corcorana/restingIAF. The following report focuses on the MATLAB implementation of the algorithm, which is dependent on the Signal Processing Toolbox™ and EEGLAB (Delorme & Makeig, 2004). The pipeline (Figure 1) relies on MATLAB’s pwelch implementation of Welch’s modified periodogram method (Welch, 1967) to derive PSD estimates. This requires the selection of a sliding window function of *x* length, which determines the frequency resolution of the analysis. (Note, alternative methods of PSD estimation could be coupled with the SGF routine, but are not explored here.) The following parameters must also be specified in order to execute the algorithm (examples of what we consider to be reasonable values are outlined in Section 2.3.4):

- *F*_*w*_, SGF frame width (longer = more smoothing; Bromba & Ziegler, 1981);
- *k*, SGF polynomial degree (higher = less smoothing/peak height loss; Press, Teukolsky, Vetterling, & Flannery, 1992);
- *W*_*α*_, the domain of the PSD searched for evidence of peak activity;
- *minP*, the minimum power value that a local maximum must exceed to qualify as a peak candidate (defined as 1 s.d. above the power estimate predicted by a regression model of the log-transformed PSD);
- *pDiff*, the minimum proportion of peak height by which the highest peak candidate within *W*_*α*_ must exceed any competitors to be assigned as the PAF;
- *cM in*, the minimum number of channel estimates necessary for computing cross-channel averages.

**Figure 1:**
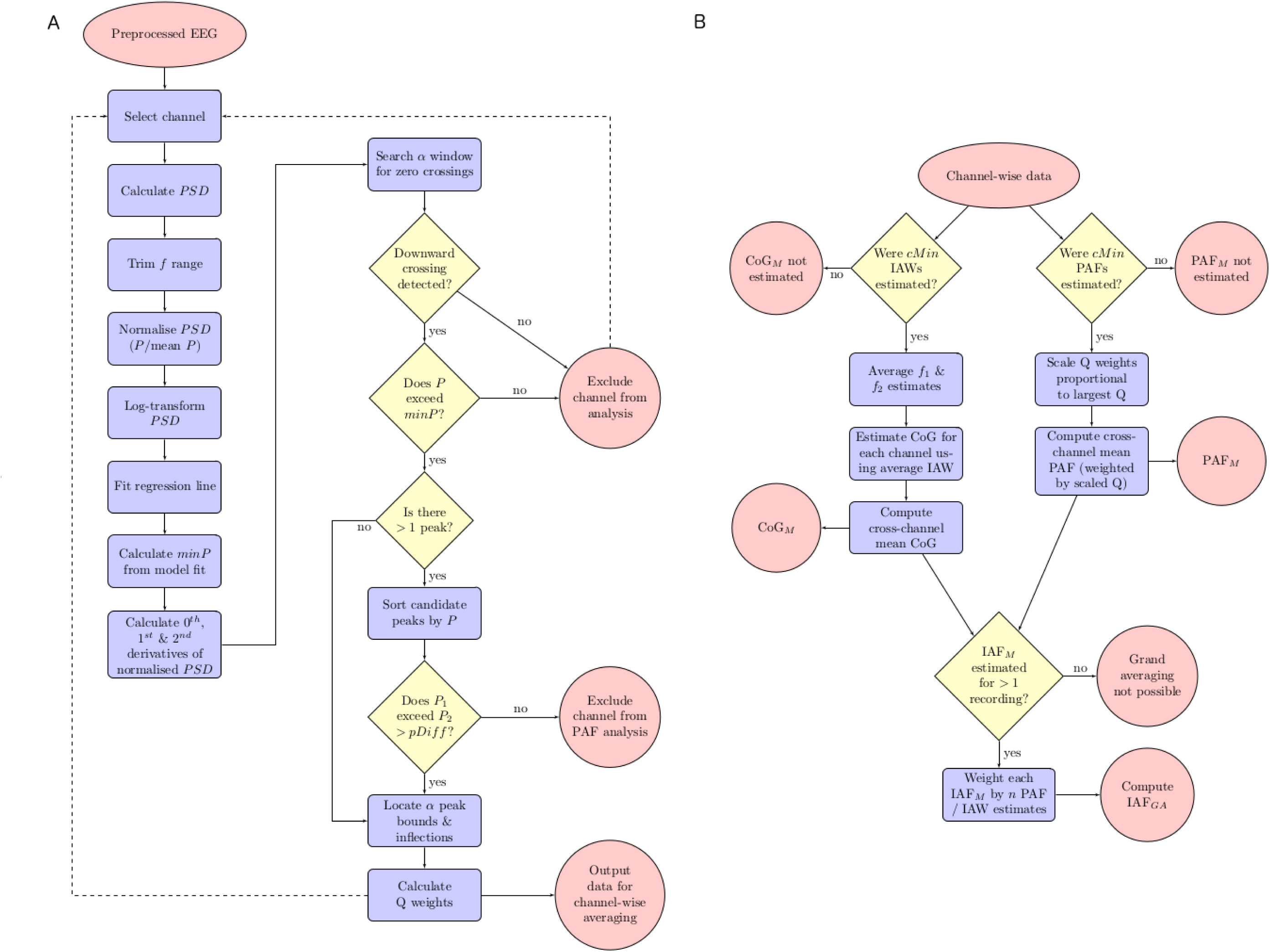
Flow diagrams summarising key steps of the analysis pipeline. (A) depicts processing of channel data, (B) depicts cross-channel averaging, assuming a sufficient number of estimates. See main text/Appendix for details. PSD: power spectral density; *f range*: frequency bins included in analysis; *P*: power estimate; *minP*: minimum power necessary to qualify as a candidate peak; *pDiff*: minimum power difference necessary to qualify as a PAF estimate; *Q weights*: quantification of relative peak quality (scaled *Q* value); *cMin*: minimum number of channel estimates required for cross-channel averaging; *IAW*: individualised alpha-band window; *f*_1_ and *f*_2_: lower and upper bounds of IAW; *PAF*_*M*_: mean PAF estimate; *CoG*_*M*_: mean CoG estimate; *IAF*_*M*_: *PAF*_*M*_ or *CoG*_*M*_; *IAF*_*GA*_: grand average PAF/CoG estimate.

Since channel spectra may be differentially contaminated by signal noise, our algorithm evaluates the relative ‘quality’ of channel-wise PAF estimates prior to cross-channel averaging. To this end, we extend the logic of the first-derivative test to extract second derivative estimates of the inflection points bounding the PAF. These points are used to define the area under the peak (normalised power units), which is then divided by the frequency span of this area. The resulting quantity (*Q value*) thus affords an indication of the relative quality of the resolved peak in terms of how well its distributional characteristics conform to the ideal of a highly powered, less variable (i.e. narrower) peak (as opposed to broader and/or shallower counterparts). Within-subject channel estimates are scaled in proportion to the peak with the highest *Q value*, and the (weighted) cross-channel average computed (hence, channels with the strongest evidence of PAF detection contribute more information to the mean estimate of the PAF). We consider this strategy (which only influences results when channel estimates fail to converge) an acceptable trade-off between loss of information (incurred by higher rates of channel exclusion) vs. loss of precision (incurred by treating all estimates as equally indicative of the estimand).

### 2.3 Empirical EEG data

#### 2.3.1 Participants

Sixty-three right-handed (Edinburgh Handedness Inventory; Oldfield, 1971), native English-speaking adults (42 females, mean age = 35 years, range = 18–74 years) with normal (or corrected-to-normal) vision and audition, and no history of psychiatric, neurological, or cognitive disorder, participated in the study. All participants provided written, informed consent, and were remunerated for their time. This study was part of a larger research project investigating EEG responses to complex, naturalistic stimuli, and was approved by the University of South Australia Human Research Ethics Committee (Application ID: 0000035576).

#### 2.3.2 Procedure

Participants were seated in a dimly-lit, sound-attenuated room for the duration of the session (2.5–3 hr). Two sets of resting-state EEG recordings were acquired approximately 90 min apart at the beginning and end of an experimental procedure. This experiment involved watching approximately 70 min of prerecorded television programming, followed by an old/new cued recall task. As per our standard laboratory protocol, both sets of resting-state recordings comprised approximately 2 min of eyes-open EEG followed by 2 min of eyes-closed EEG. Participants were instructed to sit still, relax, and avoid excessive eye movements during this time. Note, only data from the eyes-closed component of the resting-state recordings are analysed here. We favour eyes-closed resting-state data on the basis that it demonstrates (1) greater interindividual variability in alpha power (Chen, Feng, Zhao, Yin, & Wang, 2008), and (2) higher within-session reliability and test-retest stability of IAF estimates (Grandy et al., 2013b) than eyes-open data. Eyes-closed recordings may also be advantageous in reducing ocular artifact.

#### 2.3.3 EEG acquisition and preprocessing

EEG was recorded continuously from 64 cap-mounted Ag/AgCl electrodes via Scan 4.5 software for the SynAmpsRT amplifier (Compumedics^®^ Neuroscan™, Charlotte, NC, USA). The online recording was digitised at a rate of 1000 Hz, bandpass filtered (passband: 0.05–200 Hz), and referenced to the vertex electrode (AFz served as the ground electrode). Eye movements were recorded from bipolar channels above and below the left eye, and on the outer canthi of both eyes. Electrode impedances were maintained below 12.5 kΩ.

EEG data acquired from eyes-closed resting-state recordings were preprocessed in MATLAB 2015a (v8.5.0.197613). All EEG channels were imported into MATLAB via EEGLAB (v13.6.5b) and re-referenced to linked mastoids. Each dataset was then trimmed to retain only the EOG and the nine centro-posterior electrodes constituting the region of interest for this analysis: Pz, P1/2, POz, PO3/4, Oz, O1/2. These channels were subjected to zero-phase, finite impulse response highpass (passband: 1 Hz,-6 dB cutoff: 0.5 Hz) and lowpass (passband: 40 Hz,-6 dB cutoff: 45 Hz), Hamming-windowed sinc filters. Automated artifact detection routines were then applied to identify regions of channel data (segmented into 2 s epochs) that contained excessive deviations in the frequency domain (frequency range: 15–40 Hz, spectral threshold: 10 dB). Channels that exhibited an improbable signal distribution (kurtosis *z*-score > 5) were excluded from analysis. EOG channels were removed following artifact rejection, and remaining channels were downsampled to 250 Hz in preparation for spectral analysis. Datasets exceeding 120 s were trimmed to this duration in order to reduce variability in the quantity of data analysed per participant.

#### 2.3.4 IAF analysis parameters

Initial parameters for the IAF analysis were determined on the basis of preliminary testing on an independent dataset (collected as part of a separate EEG protocol). We implemented pwelch with a 1024 sample Hamming window (i.e. 4 times the sampling rate raised to the next power of 2; window overlap = 50%), yielding a frequency resolution of ~0.24 Hz. SGF and peak detection parameters were defined as follows: *F*_*w*_ = 11 (corresponding to a frequency span of ~2.69 Hz); *k* = 5; *W*_*α*_ = [7, 13 Hz]; *pDif f* = .20 (meaning that the largest peak detected within *W*_*α*_ had to be at least 20% higher than any other peak to qualify as the PAF estimate); *cMin* = 3. *minP* was automatically determined for each channel spectrum according to its distributional characteristics.

### 2.4 Simulated EEG data

#### 2.4.1 Single component simulations

As an initial proof of concept, we analysed the performance of the SGF routine in extracting target alpha frequency components embedded within noisy time series. These composite signals were created by combining a sine wave oscillating at target frequency *Fα* with a 2 min ‘pink noise’ signal (i.e. a randomly sampled signal with a frequency distribution scaled in accordance with the 1*/f* inverse power-law). SNR was manipulated by varying the length of the target signal embedded in the composite time series (e.g., for SNR = 0.5, the first half of the signal would comprise the convolution of the alpha and noise signals, whereas the second half would comprise only the noise signal).

We examined PAF estimation at SNR = 0.05, 0.10, 0.15, 0.20, 0.25, 0.30, 0.40, and 0.50, generating 1000 simulated signals for each SNR level. The target frequency was randomly sampled (with replacement) from a vector ranging from 7.5 to 12.5 in iterations of 0.1. We compared the SGF routine’s capacity to extract these target peaks with a that of a simple peak detection routine designed to locate the local maximum (LM) within *W*_*α*_. To avoid spurious estimates from suprema at the lower bound of *W*_*α*_, this routine evaluated whether the LM exceeded the power estimates of adjacent frequency bins (thus making it functionally equivalent to the first-derivative test).

#### 2.4.2 Mixture and multi-channel simulations

Next, we investigated the performance of the SGF routine under more ecologically valid spectral conditions. This involved creating alpha signals that were comprised of a set of neighbouring frequency components from different channels. We did this by sampling an ‘actual’/‘measured’ alpha frequency per channel from a truncated Gaussian distribution centered at the randomly sampled target *Fα* (selected as for the single component simulation) for each simulated (sub)component (targets chosen uniformly from the standard alpha band, as above). The tails of the Gaussian were truncated *±* 2.5 Hz from its mean/target frequency. Alpha signals were thus constructed by creating a weighted average of frequencies within this distribution; in other words, a Gaussian blur was applied to the frequency-domain signal in order to generate a mixture of alpha waves in the time domain.

Constructed alpha signals were again combined with random pink noise signals at a specified SNR. This time, each composite alpha signal was replicated 9 times, and combined with an independently sampled pink noise signal. This yielded a dataset of 9 synthetic ‘channels’, each comprised of identical alpha signals embedded within stochastically varying background noise. This enabled us to examine how our algorithm’s channel exclusion and averaging procedures performed under varying levels of SNR and peak dispersal.

As per the preliminary analysis, we compared the accuracy of SGF-generated PAF estimates against those produced by the LM procedure. For the latter, the optimisation function was applied to the mean PSD calculated for each set of simulated channel data. The simulation of broader alpha-band components also afforded the opportunity to assess the performance of the CoG estimator implemented in the SGF routine.

Finally, we repeated the multi-channel simulations using a set of alpha signals sampled via a bimodal Gaussian window. This analysis was designed to replicate troublesome empirical cases in which IAF calculation is complicated by the presence of a split-peak; either through poor resolution of a single underlying component, or where dominant activity across multiple alpha-generators results in overlapping frequency components. This analysis likewise investigated the effect of modulating the composition of the alpha signal, and the SNR of the combined time series, on IAF estimation. As the bimodal sampling window introduced the possibility of more extreme peaks (since peaks necessarily fell either side of the window centre), the span of *W*_α_ was extended to [6, 14 Hz]. This exception aside, all simulation analyses implemented the same set of parameters as described in Section 2.3.4.

## 3 Results

### 3.1 Empirical EEG data

#### 3.1.1 Global performance of the SGF routine

Post-experiment resting-state recordings were missing for 3 participants. A total of 11 channels (all from separate recordings) were excluded on the basis of excessive kurtosis. This left a total 1096 PSDs to estimate across the sample (pre = 561, post = 535). Of these, a total 944 PAF (pre = 480, post = 464) and 1003 CoG (pre = 507, post = 496) estimates were extracted. As Figure 2 indicates, the estimation routine extracted a high proportion of PAF and CoG estimates across most individuals. Two participants failed to surpass the *cMin* threshold for both recordings and were therefore excluded from the PAF analysis. Visual inspection of channel spectra confirmed the absence of any consistent alpha peak. The CoG was however estimated for one of these individuals.

**Figure 2:**
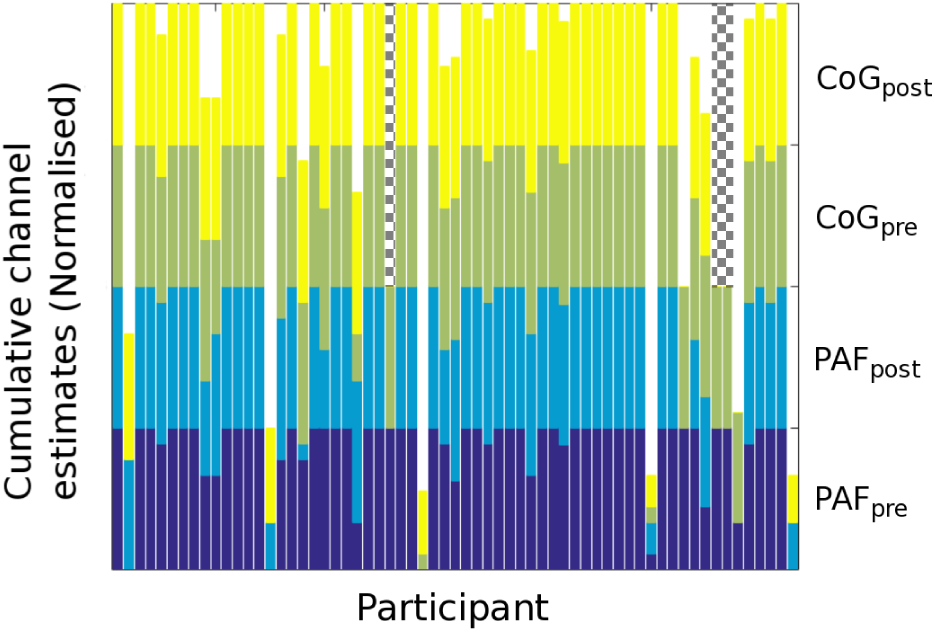
Stacked bar chart displaying number of channels from which PAF (lower half) and CoG (upper half) estimates were derived across participants. Estimates are further divided according to EEG recording (pre/post). Totals normalised to take into account excluded channels. Post-experiment data were unavailable for 3 participants (indicated by hatching).

#### 3.1.2 Statistical properties of IAF estimates

Mean IAF estimates were centred about 10 Hz, with the majority falling in the range of 9 to 11 Hz. Both estimators were similarly distributed in both sets of recordings (see Figure 3A). Intraclass correlation coefficients (ICC_3,*k*_: PAF_*M*_ = .96; CoG_*M*_ = .98) indicated that variance in PAF_*M*_ and CoG_*M*_ estimates was predominantly attributable to interindividual differences across the sample, rather than intraindividual differences between recordings (see Figure 3B). These data are therefore in accord with previous reports of the IAF’s high temporal stability (at least within the same recording session) and interindividual variability (at least in the context of eyes-closed resting-state EEG).

**Figure 3:**
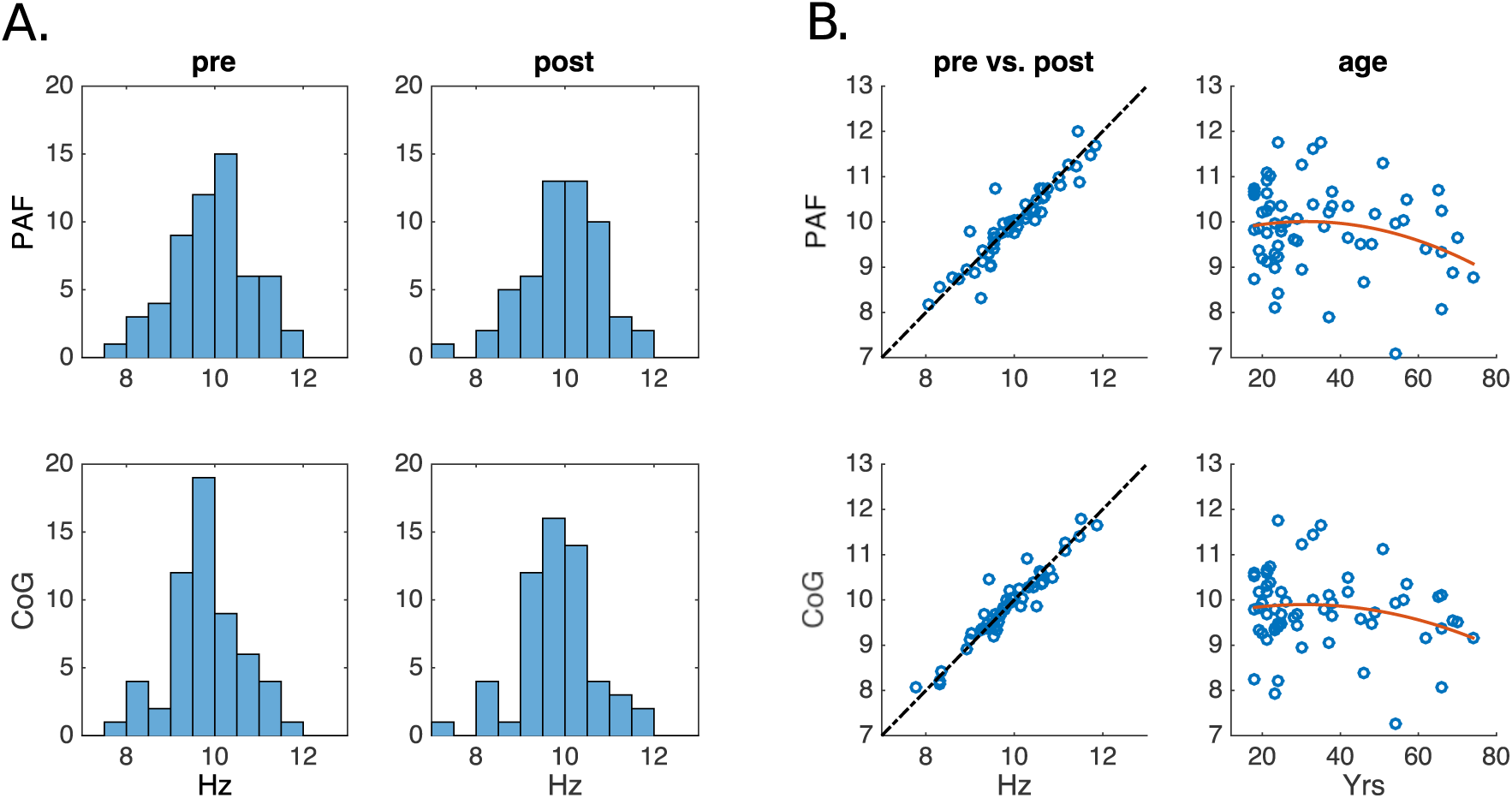
Statistical properties of PAF and CoG estimates. A: Histograms displaying distribution of mean PAF and CoG estimates across pre/post recordings. B: Scatterplots showing correlations between pre and post IAF estimates (left column), and grand-averaged IAF estimates as a function of age (right column). Broken line indicates perfect positive correlation. Red line indicates 2nd-degree polynomial fit.

Kernel density estimation of grand-averaged alpha peak and gravity estimates (PAF_*GA*_ and CoG_*GA*_, respectively) suggested that the probability density function underlying both estimators was well-characterised by a Gaussian distribution, although CoG_*GA*_ was rather more heavy-tailed. Despite this difference, PAF_*GA*_ and CoG_*GA*_ produced remarkably consistent results (ICC_3,*k*_ = .97; *R*^2^ = .90). This finding, which extends that reported in a smaller sample by Jann, Koenig, Dierks, Boesch, and Federspiel (2010), lends weight to the claim that these two estimators tap into the same fundamental oscillatory process(es).

As a final point of comparison with previous findings, we examined the relation between age and IAF (Figure 3B). Both estimators showed a similar trend towards reduced IAF as age increases beyond the fourth decade. However, this association accounted for a rather small proportion of the variance (*R*^2^ = 0.05 and 0.04 for PAF_*GA*_ and CoG_*GA*_, respectively). This is consistent with previously reported findings from much larger datasets (e.g., Chiang et al., 2011).

### 3.2 Simulated EEG data

#### 3.2.1 PAF estimator performance as a function of SNR

Preliminary analysis of synthetic EEG data focused on the number of PAF estimates extracted at each SNR level, and how well these estimates approximated the ground truth stipulated by the frequency of the alpha signal embedded in the synthetic time series. The results of this analysis are summarised in Table 1.

**Table 1:**
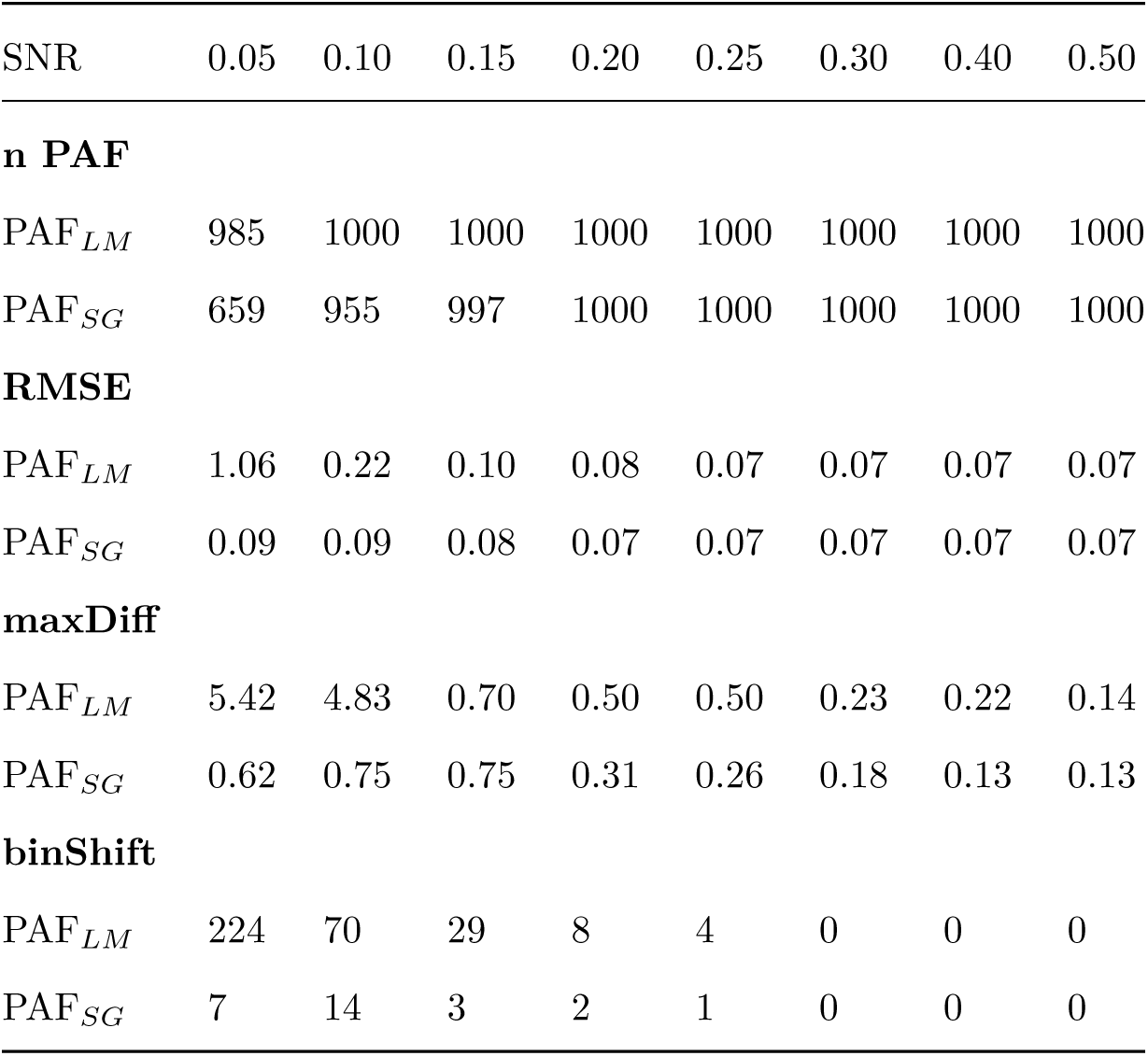
Summary statistics characterising PAF estimation as a function of estimation method and SNR. *PAF*_*LM*_: PAF estimated via the local maximum detection method; *PAF*_*SG*_: PAF estimated via the Savitzky-Golay smoothing method; *n PAF*: total number of PAF estimates extracted from 1000 simulated time series; *RMSE*: root mean squared error; *maxDiff*: maximum absolute difference between estimated and target frequency; *binShift*: number of estimates that diverged from their target frequency by > 0.24 Hz.

The SGF technique failed to extract PAF estimates for approximately one-third of simulations at SNR = 0.05, however the proportion of estimated alpha peaks rapidly approached ceiling as SNR increased beyond 0.10. Average error (*RMSE*) was generally low for all levels of SNR, suggesting that alpha peaks were consistently estimated with a high degree of accuracy when detected by the SGF analysis routine. Between 1-2% of PAF estimates in the SNR < 0.15 conditions deviated from their target frequencies by the equivalent of up to 3 frequency bins. Given the rareness of these *binShift* deviations in the higher SNR conditions, and the relatively low magnitude of such discrepancies when they did occur, it seems that the SGF technique exhibited near-optimal performance at SNR *≥* 0.30.

The LM routine returned PAF estimates for all simulated spectra; however, 15 estimates in the SNR = 0.05 condition were discarded as lower bound suprema. Even with these estimates removed, LM detection was associated with a 12-fold increase in average estimate error in the SNR = 0.05 condition as compared to the SGF method. Of the 224 estimates that were shifted by more than one frequency bin from their corresponding target frequency, 42 deviated by 1 to 2.5 Hz, while a further 56 deviated by > 2.5 Hz. All of these extreme errors constituted underestimates of the target component. The LM procedure was also markedly less accurate in the SNR = 0.10 condition, where it registered more than double the RMSE of SGF-resolved peaks. Average LM estimation error converged with that of the SGF technique in higher SNR conditions, although the magnitude of worst errors (*maxDiff*) remained elevated relative to SGF-generated PAF estimates.

To give a flavour of how smoothing may have influenced the PSD estimates generated by pwelch at each SNR level, a selection of simulated PSD functions are illustrated in Figure 4. Both techniques return identical PAF estimates at the higher SNRs. The SGF also tends to attenuate peak height, as would be expected of a smoothing procedure. The SNR = 0.30 panel reveals one instance where the application of the smoothing procedure to a reasonably blunt component results in the erroneous ascription of PAF to a neighbouring frequency bin. The advantages of the SGF technique are however thrown into relief by two scenarios where the LM estimator errs. In the SNR = 0.05 panel, the LM routine identifies a spurious fluctuation at 7.57 Hz as the PAF (*Fα* = 9.9 Hz). Here, the LM technique is disadvantaged by its inability to evaluate whether the detected LM constitutes a substantial deviation from background noise. The second scenario arises when the target component is suboptimally resolved by pwelch, resulting in either a broad structure featuring two maxima (SNR = 0.10) or a more clearly defined split-peak (SNR = 0.20). In both cases, smoothing helps to recover the shape of the peak component underlying the spectral data, thus culminating in more veridical PAF estimates than those derived via the LM method.

**Figure 4:**
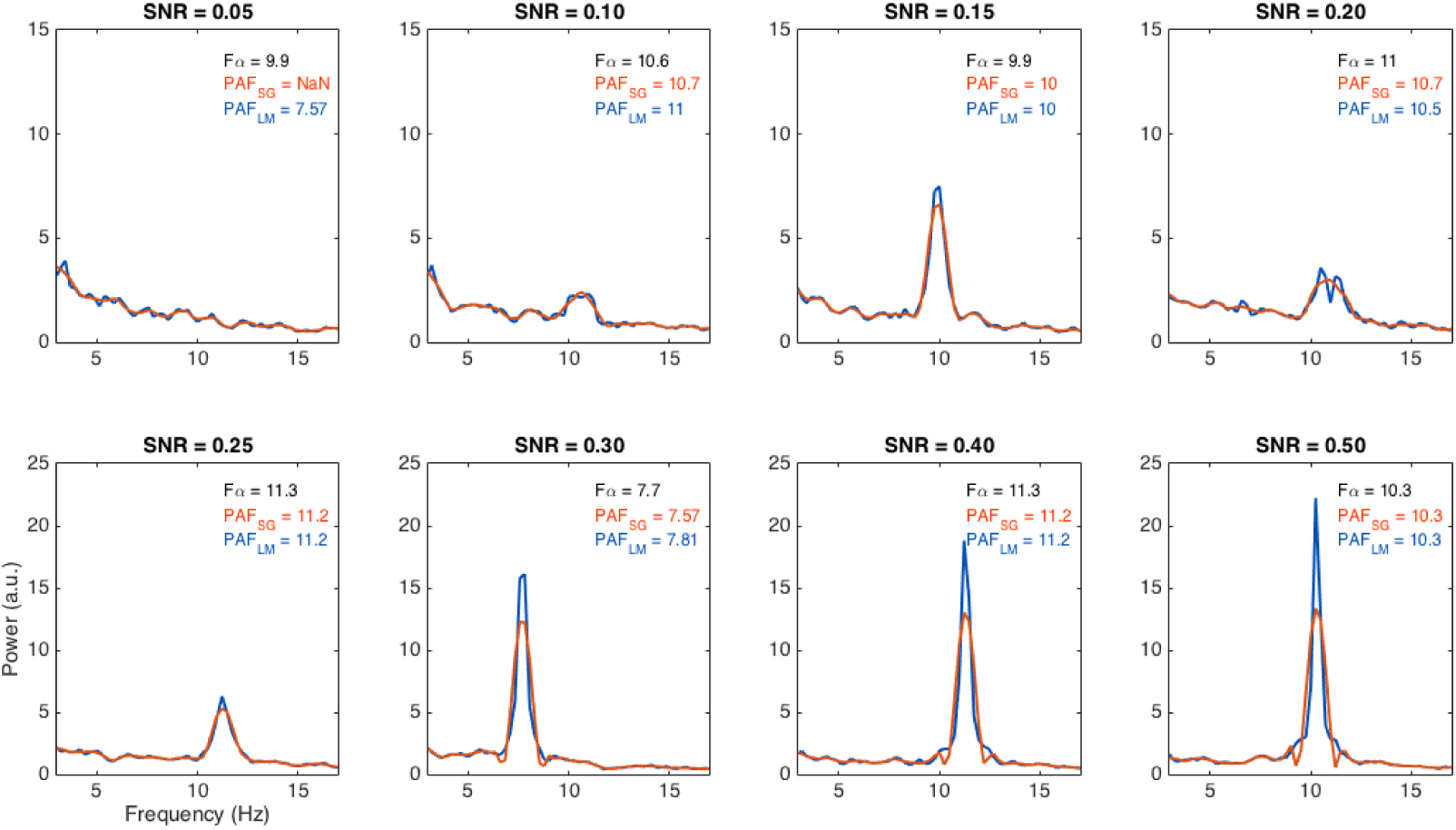
Channel spectra randomly sampled from each SNR condition. Blue functions represent PSD estimates generated by pwelch. Red functions indicate effect of smoothing these estimates with the Savitzky-Golay filter (SGF). *Fα*: Target alpha component frequency; *PAF*_*SG*_ and *PAF*_*LM*_: Estimates of *Fα* rendered by the SGF and local maximum methods, respectively. *a.u.*: Arbitrary unit; *NaN*: No estimate returned.

In sum, this preliminary analysis provides compelling evidence that the SGF method generally furnishes accurate estimates of the PAF when a singular alpha component is present within the PSD. Such accuracy is maintained even at relatively low SNR levels, although the extraction of low-powered peaks amidst background noise becomes more challenging when SNR drops below 0.15. The more conservative nature of the SGF method (as compared to LM detection) in the context of low SNR may however be advantageous in protecting against inaccurate PAF estimates issuing from spurious background fluctuations.

#### 3.2.2 Multi-channel dataset simulations

Given that the PAF estimators approached ceiling performance at moderate levels of SNR in the previous analysis, we limited multi-channel simulations to a low (0.15) and a moderate (0.40) SNR condition. A total of 100 datasets, each comprising 9 synthetic EEG channels, were simulated for each level of alpha component dispersal in both SNR conditions (yielding a total 5400 PSD estimates). The results of this analysis are summarised in Figure 5 and Table 2.

**Figure 5:**
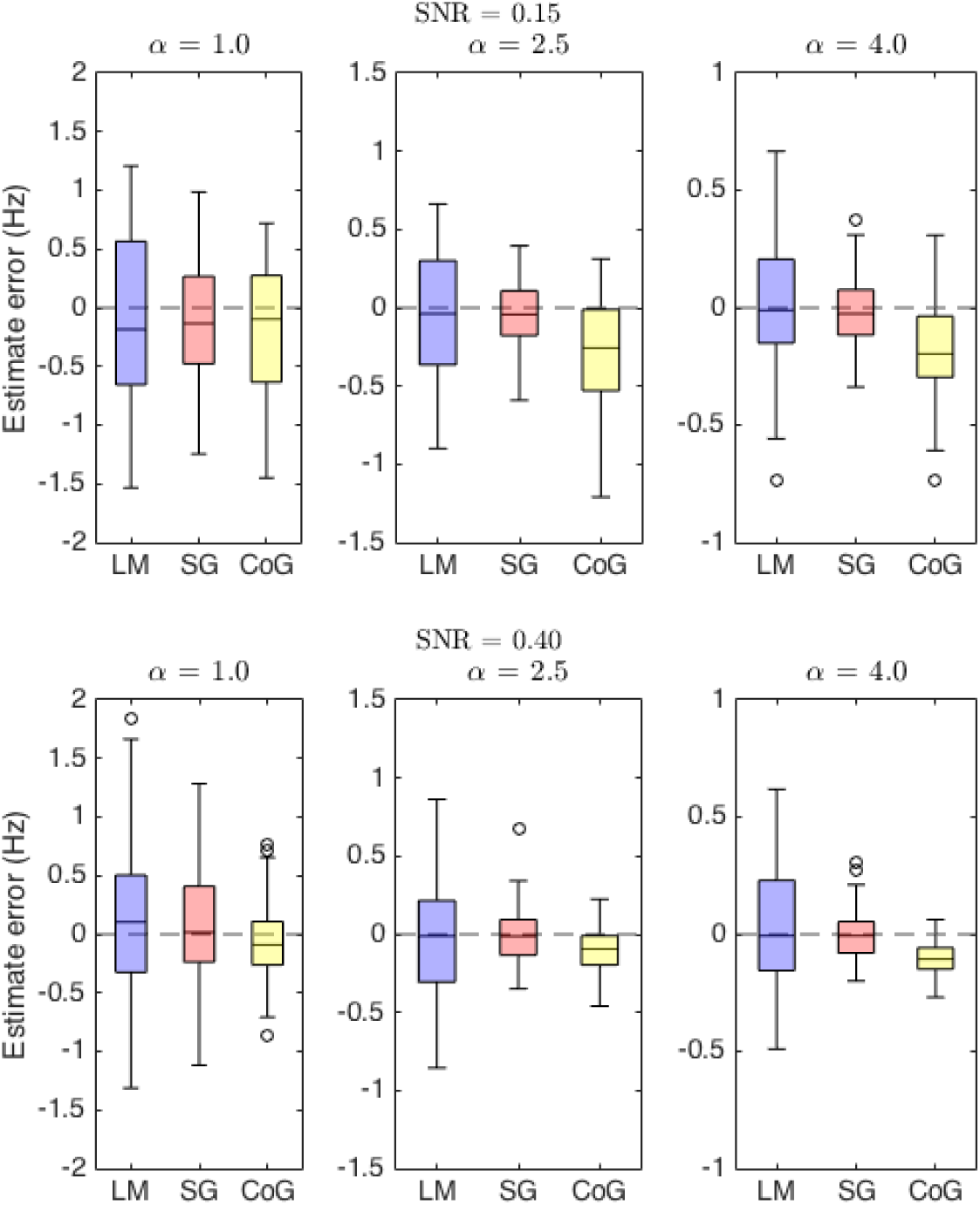
Box plots summarising spread of estimator error across simulation conditions. Centre marks indicate median error, edges indicate interquartile range (IQR), whiskers indicate approximately 1.5 × IQR. Zero estimate error (broken horizontal line) corresponds to extraction of the target alpha peak frequency. Negative error indicates underestimation of the target frequency, positive error indicates overestimation. Dispersal of the target alpha component broadest in the left column (*α* = 1.0) and narrowest in the right (*α* = 4.0). *LM* and *SG*: PAF estimated via the Local Maximum and Savitzky-Golay routines, respectively. *CoG*: CoG estimated via the Savitzky-Golay routine. Y-axis scaling varied across *α* levels to aid visualisation.

**Table 2:** Estimator performance as a function of SNR and alpha component distribution (*α* = 1.0 corresponds to a broad peak, *α*= 4.0 a narrow peak). *PAF*_*LM*_: Local maximum PAF estimator; *PAF*_*SG*_: Savitzky-Golay filter (SGF) PAF estimator; *CoG*: SGF CoG estimator; *RMSE*: root mean squared error; *maxDiff*: maximum absolute difference between estimated and target frequency; *% Dev*: percentage of estimates that diverged from the target frequency by > 0.5 Hz; *n chans*: median (s.d.) number of channels furnishing PAF/IAW estimates per simulated dataset.

| SNR | 0.15 |  |  | 0.40 |  |  |
| --- | --- | --- | --- | --- | --- | --- |
| $\alpha$ | 1.0 | 2.5 | 4.0 | 1.0 | 2.5 | 4.0 |
| **RMSE** |  |  |  |  |  |  |
| *PAF<sub>LM</sub>* | 0.72 | 0.38 | 0.30 | 0.63 | 0.38 | 0.26 |
| *PAF<sub>SG</sub>* | 0.47 | 0.21 | 0.15 | 0.48 | 0.17 | 0.10 |
| *CoG* | 0.57 | 0.45 | 0.27 | 0.34 | 0.16 | 0.12 |
| **maxDiff** |  |  |  |  |  |  |
| *PAF<sub>LM</sub>* | 1.53 | 0.90 | 0.73 | 1.84 | 0.86 | 0.62 |
| *PAF<sub>SG</sub>* | 1.24 | 0.59 | 0.38 | 1.29 | 0.68 | 0.31 |
| *CoG* | 1.45 | 1.21 | 0.73 | 0.86 | 0.46 | 0.27 |
| **% Dev** |  |  |  |  |  |  |
| *PAF<sub>LM</sub>* | 63 | 17 | 14 | 42 | 22 | 3 |
| *PAF<sub>SG</sub>* | 30 | 2 | 0 | 33 | 1 | 0 |
| *CoG* | 42 | 30 | 7 | 18 | 0 | 0 |
| **n chans (s.d.)** |  |  |  |  |  |  |
| *PAF<sub>SG</sub>* | 5 (1.81) | 6 (1.53) | 8 (1.23) | 5 (1.52) | 7 (1.27) | 9 (0.70) |
| *CoG* | 9 (0.79) | 9 (0.36) | 9 (0.10) | 9 (0) | 9 (0) | 9 (0) |

Across all Distribution × SNR conditions, the SGF routine failed to generate average PAF estimates for 11 datasets. Eight of these instances occurred in the low SNR condition (7 *α* = 1.0; 1 *α* = 2.5), while the remainder occurred when attempting to recover broad component structures (*α* = 1.0) in the moderate SNR condition. By contrast, both the LM and the CoG estimators rendered estimates for all 600 simulated datasets.

All three estimators demonstrated consistent reductions in error as alpha component dispersal diminished (i.e. as target peaks became narrower). This finding is congruent with the intuition that, irrespective of SNR, recovery of broader component structures poses a greater challenge for automated estimation procedures than the recovery of narrower, sharper peaks. Further, there was some indication of a Distribution *×* SNR interaction effect, such that error indices for a given *α* level were more elevated in the low (as compared to the moderate) SNR condition. Although this effect was somewhat marginal (and not entirely consistent) for the PAF estimators, it was more clearly apparent for the CoG estimator. These general trends (i.e. improved estimation accuracy with decreased component dispersal and increased SNR) were mirrored by both the average (median) number of channels that contributed to PAF_*SG*_ estimation, and the degree of variability (s.d.) in the number of channels retained by the SGF procedure for each set of simulations. This is to say that a higher proportion of channels rendered PAF estimates as SNR increased and peak dispersal decreased, while volatility in the number of channels selected for mean PAF/IAW estimation correspondingly declined.

As per the single component analysis, PAF estimates from low SNR simulations were more accurate on average when estimated with the SGF procedure. Unlike the prior analysis, however, the RMSE of PAF_*LM*_ failed to converge with that of PAF_*SG*_ in the moderate SNR condition (indeed, RMSE of the former was more than double that of the latter for both intermediate and narrow peak estimates). The magnitude of worst estimate errors (*maxDiff*) was likewise consistently elevated for PAF_*LM*_ as compared to PAF_*SG*_-generated estimates. Perhaps most notably, PAF_*LM*_ produced considerably more estimate errors in excess of *±* 0.5 Hz than PAF_*SG*_ (27% vs. 11%). This contrast was most stark at *α ≥* 2.5, where the error rate associated with PAF_*LM*_ was 14% (compared to < 1% for PAF_*SG*_).

Comparison of SGF-generated estimates of PAF and CoG discloses an interesting interaction between estimator performance and SNR. While the PAF estimator resulted in diminished RMSEs, lower maximal deviations, and fewer estimation errors *±* 0.5 Hz in the low SNR simulations, this pattern was inverted (with the exception of one RMSE value) in the moderate SNR condition. This latter result provides encouraging evidence in favour of our method’s capacity to accurately localise the beginning and end of the IAW (at least when the embedded alpha signal is not too weak). Interestingly, even though the CoG performed less consistently when SNR was low, it still tended to be more reliable than the PAF_*LM*_ estimator. For instance, the CoG method resulted in a 16% reduction in substantial estimate errors compared to the LM method. While CoG may therefore be more susceptible to bias than its PAF_*SG*_ counterpart when channel spectra contain relatively high degrees of background noise, it may still offer tangible advantages over LM-based peak detection strategies.

#### 3.2.3 Split-peak simulations

Finally, we repeated the multi-channel dataset simulations with composite signals constructed using a bimodal sampling window. This window comprised two overlapping Gaussians (*α* = 2.5), the right-most of which was scaled equal to, 0.25, or 0.50 times larger than its counterpart. The frequency offset between the two Gaussian peaks was equivalent to 1.6 Hz. The results of this analysis are summarised in Figure 6 and Table 3.

**Figure 6:**
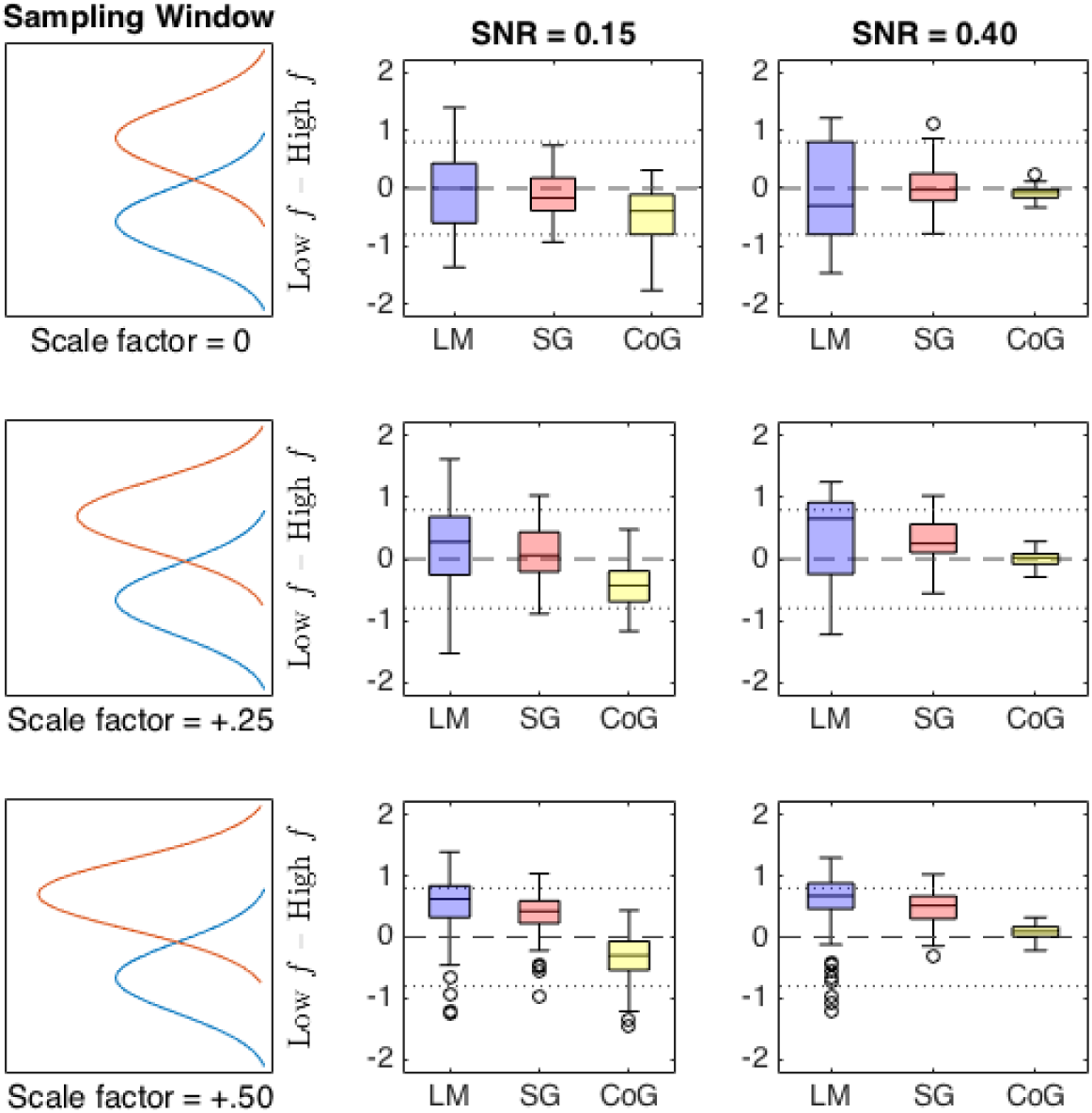
Box plots summarising spread of estimate deviation from the centre frequency of the sampling window. Centre marks indicate median deviation, edges indicate interquartile range (IQR), whiskers indicate approximately 1.5 × IQR. Zero deviation (broken horizontal line) corresponds to estimating the midpoint between the two components. Peak locations indicated by dotted horizontal lines. *Left column*: Schematic of the sampling window used to construct composite alpha signals simulated in corrosponding row. The discrepancy between simulated peaks (higher relative to lower frequency bins) ranges from 0 (top row) to +0.50 (bottom row). *LM* and *SG*: Local Maximum and Savitzky-Golay PAF estimates, respectively. *CoG*: Savitzky-Golay CoG estimates.

**Table 3:** Estimator performance as a function of SNR and relative weighting of bimodal peaks. Right-most Gaussian function was either 0, 0.25, or 0.50 times larger than the left (*PeakDiff*). *PAF*_*LM*_: Local maximum PAF estimator; *PAF*_*SG*_: Savitzky-Golay filter (SGF) PAF estimator; *CoG*: SGF CoG estimator; *RMSE*: root mean squared error (relative to centre frequency of sampled components); *maxDiff*: maximum absolute difference between estimates and centre frequency of sampled components; *n chans*: median (s.d.) number of channels furnishing PAF/IAW estimates per dataset.

| SNR | 0.15 |  |  | 0.40 |  |  |
| --- | --- | --- | --- | --- | --- | --- |
| *PeakDiff* | 0 | +0.25 | +0.50 | 0 | +0.25 | +0.50 |
| **RMSE** |  |  |  |  |  |  |
| *PAF<sub>LM</sub>* | 0.69 | 0.69 | 0.75 | 0.84 | 0.79 | 0.76 |
| *PAF<sub>SG</sub>* | 0.40 | 0.44 | 0.51 | 0.38 | 0.45 | 0.55 |
| CoG | 0.62 | 0.56 | 0.51 | 0.14 | 0.12 | 0.15 |
| **maxDiff** |  |  |  |  |  |  |
| *PAF<sub>LM</sub>* | 1.40 | 1.62 | 1.40 | 1.47 | 1.25 | 1.30 |
| *PAF<sub>SG</sub>* | 0.93 | 1.03 | 1.04 | 1.10 | 1.03 | 1.03 |
| CoG | 1.77 | 1.17 | 1.47 | 0.33 | 0.29 | 0.32 |
| **n chans (s.d.)** |  |  |  |  |  |  |
| *PAF<sub>SG</sub>* | 4 (1.52) | 5 (1.49) | 5 (1.68) | 5 (1.36) | 6 (1.64) | 6 (1.31) |
| CoG | 9 (0.46) | 9 (0.48) | 9 (0.51) | 9 (0) | 9 (0) | 9 (0) |

PAF_*SG*_ failed to find evidence of a distinct peak in 11% of low SNR datasets (Equal = 14, +0.25 = 7, +0.50 = 11), and 2% of moderate SNR datasets (Equal = 3, +0.25 = 4, +0.50 = 0). Median number of channel PAF estimates was also reduced as compared to the corresponding SNR conditions in the single-peak, multi-channel simulations. As per the single peak, multi-channel simulations, both PAF_*LM*_ and CoG returned estimates for all 600 simulated datasets.

Across all conditions, PAF_*LM*_ returned more variable and extreme results than PAF_*SG*_; although interpretation of this observation is complicated by the presence of a (somewhat) dominant peak in the +0.25 and +0.50 conditions. As both SNR and peak difference increase, PAF_*LM*_ shows stronger migration towards the higher frequency peak than either of the SGF estimators, although note that it is still more prone to erroneously ascribing the PAF to the secondary (lower frequency) peak. On the other hand, PAF_*SG*_ is less liable to spurious fluctuations in the PSD, tending instead to curb PAF estimation towards the centre mass of the double component. This might suggest that marginal local maxima are absorbed within the recapitulation of a broader peak structure as a consequence of spectral smoothing. As SNR and peak inequality increase, PAF_*SG*_estimates cluster in closer proximity to the dominant peak. This then explains why RMSE increases relative to the centre frequency: as SNR improves and the split-peak becomes more asymmetrical (and hence, one peak more dominant over its competitor), more evidence accrues in favour of an underlying PAF.

The CoG estimator an intermediate level of variability compared to the PAF estimators under low SNR conditions, but is markedly less variable under moderate SNR conditions. The box plots in Figure 6 also indicate that the CoG estimator performed similarly across the different degrees of peak inequality within each SNR level. Irrespective of peak scaling, CoG estimates were substantially more precise when SNR = 0.40. Indeed, compared to the other two estimators, CoG is both remarkably stable and closely centred around the centre frequency of the window function. As such, this finding provides compelling evidence that our implementation of the CoG estimator renders an accurate summary of the underlying alpha component distribution.

## 4 Discussion

We have proposed a novel method for estimating the two most prevalent indices of individual alpha frequency (IAF) in the literature. This method pairs a common approach to the automated detection of local maxima (i.e. searching for first derivative zero crossings) with a well established method of resolving spectral peaks (i.e. Savitzky-Golay filtering) to derive an estimate of peak alpha frequency (PAF). It also extends the logic of the first-derivative test to estimate the bounds of the alpha peak component, thus enabling calculation of the alpha-band centre of gravity (CoG). Like other automated curve-fitting algorithms reported in the literature (e.g., Chiang et al., 2008; Lodder & Putten, 2011), this method addresses key limitations of visual PSD analysis (e.g., proneness to subjective bias, inefficiency, and poor replicability), while improving upon alternative automated approaches that may be prone to various artifacts (e.g., failure to differentiate a single dominant peak from competing spectral peaks or spurious fluctuations, reliance on alpha-band reactivity). Unlike these algorithms, however, our method is openly accessible and easy to integrate within existing MATLAB and Python-based analysis pipelines.

Our results demonstrate that the SGF technique can extract a high proportion of IAF estimates from an empirical dataset, and that the sample-wide properties of these estimates (intraindividual stability, interindividual variance, etc) are consonant with prior reports in the literature. Furthermore, application of the technique to simulated datasets verified its ability to render accurate estimates of peak location, even under highly degraded SNR conditions. When extended to more complex simulations, the SGF technique was shown to recover target values with greater precision than an alternative peak detection method. We begin by considering the key findings of our analyses, before reflecting on present limitations and potential directions for future research.

### 4.1 Estimation of IAFs from an empirical EEG dataset

Savitzky-Golay filtering of pwelch-generated PSD functions resulted in the extraction of a rather impressive number of IAF estimates from a moderate-sized dataset. This suggests our technique offers substantive benefits in terms of data retention in comparison to subjective analysis, which can result in high rates of attrition if dominant peaks cannot be confidently distinguished from background noise (e.g. Bornkessel-Schlesewsky et al., 2015). We note also that our SGF method furnished a higher proportion of PAF estimates than that produced by the Gaussian curve-fitting procedure implemented by Haegens and colleagues (2014). It may be the case that our non-parametric approach, which attempts to smooth the PSD rather than fit a specified function to it, retains more data by virtue of its capacity to accommodate a broader range of alpha-band distributions.

By the same token, it is reassuring that neither of the two cases in which the technique failed to extract PAF estimates demonstrated compelling evidence of any concerted alpha peak activity on visual inspection of their respective PSD plots. It is also worth pointing out that the diverse age range of participants within this study is likely to have posed a nontrivial challenge to any automated alpha-band quantification routine, given the typically reported changes in both spectral power and distribution associated with older adulthood (e.g., Dustman, Shearer, & Emmerson, 1999). That our technique was able to extract estimates for the vast majority of sampled individuals, and that it did so using a fixed set of parameters defined a priori on the basis of preliminary testing in an independent dataset, speaks to its capacity to derive resting-state IAF estimates across a broad spectrum of the healthy population.

Comparison of grand-averaged PAF and CoG estimates revealed a high degree of intercorrelation, despite certain differences in their distribution. Although this might prompt concerns of redundancy, we interpret this finding positively: the CoG seems to tap into a similar underlying neural process (or set of processes) as the PAF. Although not necessary in the present analysis on account of the high proportion of PAFs that were extracted across participants, this finding suggests that the CoG estimator might warrant deployment as an alternative marker of IAF in cases where the PAF cannot be obtained. In any case, given the dearth of research directly comparing these two measures (most IAF-related research involves some variant of PAF, perhaps on account of the additional complexities involved in calculating the CoG), we suggest it would be informative if investigators adopted the policy of reporting both indices in parallel. Should it be the case that PAF and CoG track one another almost identically, then only one of these measures need be selected for the remaining analysis (see for e.g., Jann et al., 2010). However, if it turns out that PAF and CoG diverge under certain circumstances, delineating such cases might help improve our understanding of the IAF (and alpha-band dynamics more generally). It is of course a notable advantage of our method that it enables investigators to rapidly derive sample-wide estimates of PAF and CoG simultaneously, thus furnishing a convenient means of estimator comparison. To the best of our knowledge, no previously reported automated technique provides this functionality.

### 4.2 Estimation of simulated IAFs

Our preliminary simulation analyses indicated that the SGF technique approached an optimal level of performance when 2 min synthetic signals featured approximately 36 s of alpha-band oscillations (SNR = 0.30). Indeed, the peak detection routine performed reasonably well when signals contained as little as 12 s of alpha-band activity, with fewer than 6% of simulated alpha components undetected or erroneously estimated by more than one frequency bin.

Interestingly, our analysis shows that less sophisticated approaches to peak estimation can result in substantial error at comparably low levels of SNR. It is likely that most of these inaccurate estimates derived from spurious local maxima occurring due to fluctuations in background spectral activity. Indeed, the LM method’s propensity to underestimate PAF in low SNR conditions supports this interpretation, since the inverse power-law (which is not generally taken into account by LM detection methods) increases the probability of spurious local maxima at lower frequencies within the search window. Such artifacts are undesirable not only for the obvious reason that they introduce additional noise into IAF-related analyses, but also insofar as such errors diminish confidence in automated analysis methods (after all, such errors would presumably have been avoided had spectral data been subjected to visual inspection). Indeed, we consider it preferable that an automated peak detection routine should reject spectra showing inconclusive evidence of any concerted alpha-band activity, rather than generate highly deviant estimates of the underlying (albeit weak) signal. It is a strength of the SGF technique, then, that it applies more stringent criteria in the evaluation of candidate peaks.

In addition to demonstrating that the SGF technique performs consistently well in low-to-moderate SNR conditions, our analysis also confirmed that the application of this smoothing procedure did not cause excessive distortion of PAF estimates. Furthermore, our analysis highlighted that discrete Fourier analysis methods (such as Welch’s modified periodogram) might precipitate artifactual split-peaks, and that such cases can be ameliorated by means of a smoothing procedure. Consequently, the single component simulation analysis stands as a basic proof of concept that the SGF method is capable of (1) extracting a high proportion of underlying peak frequencies without introducing systematic bias, and (2) improving upon existing techniques of peak resolution and estimation, thus helping to maximise the number of IAF estimates that can be extracted from a given dataset. We acknowledge however that the estimation of sharply defined, single frequency alpha components may well be unrepresentative of genuine electrophysiological data in many contexts. While it is encouraging then that the SGF technique performed well under these reasonably favourable conditions, it was necessary to demonstrate its capabilities when confronted with more complex, ecologically valid signals.

The multi-channel simulation analyses were designed to be more faithful to empirical resting-state EEG data, in as much as each target signal comprised a range of alpha components embedded within a variety of nonidentical (but highly correlated) time series. These simulations also enabled us to examine the performance characteristics of the SGF routine’s CoG estimator, which was expected to closely approximate the PAF in the context of Gaussian-distributed alpha components. The critical finding across all simulation conditions was that the SGF technique rendered PAF and CoG estimates that almost always improved upon LM-derived PAF estimates from averaged channel spectra. This finding held irrespective of whether estimator deficits were quantified in terms of the average error across simulated datasets, magnitude of worst (i.e. most deviant) estimate errors, or percentage of estimates in the dataset that deviated from the ground truth by more than *±* 0.5 Hz (a threshold previously used by Lodder and Putten, 2011, to evaluate the performance of their peak detection algorithm).

Leaving aside the superiority of the SGF over the LM detection routine, one might still raise the concern that its performance falls somewhat short when applied to broadly-dispersed alpha component structures. Indeed, the RMSE of the PAF estimator in both SNR conditions of the single-peak analysis approaches the *±* 0.5 Hz threshold demarcating substantial estimate deviation, while the CoG exceeds this limit when SNR is low. Correspondingly, low-*α* multi-channel simulations returned a much higher proportion of estimates exceeding the *±* 0.5 Hz error threshold (as compared to simulations involving higher *α* levels), especially in the case of the PAF estimator. It ought to be borne in mind, however, that all simulation analyses were performed using SGF parameters identical to those used in the empirical analysis. This is pertinent because it is likely that the filter frame width (*F*_*w*_ = 11) was suboptimally narrow for the purpose of smoothing such broad peak structures. Indeed, post hoc analysis (not reported) revealed that simply doubling the length of the filter frame can halve the number of simulations that failed to produce PAF estimates, as well as reducing substantial estimate deviation by one third under moderate SNR conditions. Corresponding improvements were not realised however in the context of low SNR; hence, the recovery of broadly dispersed, relatively weak alpha signals remains technically challenging.

Of the three IAF estimators examined in these simulations, the CoG was most sensitive to manipulation of the SNR. That low SNR simulations should inflict notable performance decrements is hardly surprising, however, given that CoG calculation depends upon the spectral characteristics of the entire (individualised) alpha-band interval across all available channels. Not only does low SNR pose nontrivial difficulties in defining the bounds of the alpha interval (thus threatening to introduce noise by either including extraneous data from beyond the alpha interval, or excluding portions of the alpha band from analysis), the relative weakness of the alpha signal means that a higher proportion of background noise contributes to CoG calculation. This scenario may be compounded by the fact that the traditional method of computing CoG estimates averages across all available channels, not just those that contributed to calculation of the IAW (although note that the average number of channels selected to infer this bandwidth remained high even in the doubly challenging conditions posed by the low SNR × broad component dispersal combination of the single-peak analysis). It might be the case then that the central tendency-like properties of the CoG, which may have underpinned its strong performance in the moderate SNR simulations (where, of the three estimators, it was the least prone to substantial estimate deviation), render it more vulnerable to error when substantive alpha-band activity is relatively sparse. Consequently, it could be useful to investigate whether the performance of the CoG estimator in relatively noisy conditions can be augmented through the development of more robust methods of calculation.

Taking the results of the single-and split-peak simulations together, it is tempting to conclude that the PAF estimator outperforms its CoG counterpart in the former scenario, while the opposite is true in the latter. Even under relatively favourable spectral conditions, the CoG estimator tended to underestimate the target frequency in the single-peak simulations. Indeed, CoG estimates increasingly deviated from the centre frequency of the target component as the latter became narrower, which seems counterintuitive if such peaks ought to be less difficult to resolve and parameterise. We suggest however that this tendency derived from the skewness introduced into the Gaussian-distributed target components when they were combined with the pink noise signal. This observation thus reinforces the point that PAF and CoG estimators summarise different features of the spectral distribution, and that they need not always converge. Analysis of the split-peak simulations suggests however that the SGF method may still be somewhat prone to PAF estimate distortion when the underlying pwelch routine fails to consistently resolve dual subcomponents across channel spectra. This finding suggests a more stringent *cM in* criterion might be advisable to avoid PAF estimates that might in fact reflect a more CoG-like average across channels that, due to random noise fluctuations, resolve only one of two (or more) underlying subcomponents. In our view, the fact that the SGF approach to PAF estimation does not fully eliminate the methodological and conceptual challenges posed by split-peaks is not so much an intrinsic shortcoming of our technique in particular, but reflects rather a problematic attribute of the PAF in general. These data thus lend weight to the argument that the CoG, insofar as it avoids these difficulties, might be preferable to the PAF.

### 4.3 Limitations and future developments

We aimed to design an accessible, fast, automated routine that calculates reliable PAF and CoG estimates from posterior channel EEG data recorded during short periods of relaxed, eyes-closed wakefulness. Although limited in its current scope, we believe that the programme could be adapted for application across a broader range of empirical contexts (e.g., quantifying spectral dynamics across various frequency bands during task-related activity; quantifying peak characteristics across different topographical regions). It may prove more challenging, however, to accurately resolve estimates of IAF under conditions that are less conducive to the manifestation of a dominant alpha peak (or indeed, in populations known to manifest spectral characteristics that differ from those of neurotypical adults). Further research would therefore be required to establish the utility of the SGF technique for applications beyond the rather circumscribed conditions examined here.

One aspect of performance that was not investigated in our analysis was whether the accuracy and precision of IAF estimates depend upon the method used to derive underlying PSD estimates. In its present implementation, our algorithm relies upon Welch’s method to estimate the PSD that is subjected to the SGF’s smoothing and differentiation operations. It may therefore be worthwhile to investigate whether alternative methods of PSD estimation (e.g., the multitaper method, continuous wavelet transform) can be exploited in conjunction with the SGF technique in order to further improve IAF estimation.

Another possible avenue for optimising the performance characteristics of the SGF routine would be to develop a function that automatically adapts the *F*_*w*_ (filter width) and *k* (polynomial degree) parameters in accordance with the approximate span of the dominant frequency component located within the search window *W*_*α*_. This would involve implementing an iterative fitting process, where the empirical features of the alpha-band component are initially parameterised in order to scale *F*_*w*_ and *k*. Once these parameters have been determined for the data at hand, smoothing and estimation procedures would proceed as described above.

Finally, it would be desirable to create a package that incorporates the MATLAB implementation of the SGF routine within the EEGLAB graphical user interface. Not only would this help to make the procedure accessible to the broadest possible range of EEGLAB users, it would also provide a convenient platform for integrating visualisations of the spectral analysis that may (for instance) assist in the diagnosis of suboptimal parameter settings. We intend to explore a number of these possibilities in future work.

## 5 Conclusion

We have developed a free, open-source programme for automatically estimating individual alpha frequency in resting-state EEG data. This programme has been shown to perform more accurately than a simpler automated peak detection routine, and may return a higher proportion of empirical IAF estimates than techniques relying on parametric curve-fitting procedures. Furthermore, this method is not dependent on phasic changes in alpha-band reactivity, which may produce biased IAF estimates. In addition to its obvious advantages from the perspective of replicability and efficiency, our simulations indicate that this method could help to improve the accuracy and precision of future IAF-related research. This technique may also open up new lines of methodological inquiry, insofar as it facilitates the direct comparison of two prevalent indices of IAF that have for the most part been investigated in isolation of one another.

## Acknowledgements

We thank Jessica Gysin-Webster and Daniel A. Rogers for their assistance with data collection.

